# Using time-resolved auditory frequency tagging to capture self-preferential processing of the own (vs other) name

**DOI:** 10.1101/2025.09.03.673961

**Authors:** Danna Oomen, Emiel Cracco, Annabel D. Nijhof, Jan R. Wiersema

## Abstract

The own name is an important, salient self-related stimulus and ostensive cue. Prioritized processing of one’s own name plays a key role in attention, self-awareness, and social and cognitive development. Previous studies using event-related potentials (ERPs) have reported a self-preferential effect for the own name, characterized by enhanced neural responses compared to the names of (close) others. In this study, we aimed to develop and validate an EEG auditory frequency-tagging task to measure self-preferential processing of the own name, leveraging the superior signal-to-noise ratio of this technique, which is highly relevant for future studies on infants and clinical populations. To this end, we ran two separate studies (dichotic: *n*_1_ = 31, non-dichotic: *n_2_* = 32). In contrast to previous ERP research, we did not find evidence of a self-preferential effect. We reasoned that collapsing temporal information may have led to the failure to capture the effect, which ERP research has shown to emerge at a late stage of processing, and that this effect may have been overshadowed by early brain responses. To address this issue, we applied a novel approach that we refer to as the *time-resolved frequency tagging approach*, which incorporates knowledge of the effect in the temporal domain. This did result in a clear self-preferential effect of the own name. Hence, we were able to develop and validate an EEG frequency-tagging task to measure the self-preferential effect of the own name. Our approach can also be used in future EEG frequency-tagging studies investigating other complex cognitive processes.

---

The own name is a highly salient stimulus, as it is a self-related stimulus (i.e. it has high personal relevance) and an ostensive cue (i.e. used by others to initiate social interaction). The salience of the own name can be nicely illustrated through the cocktail party effect (Cherry, 1953). This effect refers to the ability to focus one’s attention on a specific task or auditory stimulus, such as a conversation, while filtering out competing sounds in a noisy environment, such as at a cocktail party. However, even though people are quite good at filtering out noise, hearing one’s own name usually captures attention even when it is irrelevant to the conversation at hand. This phenomenon has been demonstrated in controlled behavioural experiments. Moray (1959) first demonstrated this with a dichotic listening task in which participants were presented with two different auditory stimuli simultaneously, directed into different ears via headphones. When repeating the stimuli presented to one of the ears out loud (referred to as ‘shadowing’), subjects were generally unable to report the content presented to the other ear. Their own name was, however, found to be an exception to the rule, highlighting the unique salience of this stimulus. This attentional advantage has been replicated in subsequent studies (Conway et al., 2011; Wood & Cowan, 1995) and has also been demonstrated for the written own name. These studies have shown that the written own name is resistant to the attentional blink and repetition blindness (Arnell et al., 2010; Nijhof et al., 2021; Shapiro et al., 1997), that presenting stimuli together with the written own name can produce a memory advantage (Kim et al., 2019), and that the written own name facilitates faster reaction times even when presented subliminally (Alexopoulos et al., 2012).

Besides behavioural methods, neuroscientific methods have been used to investigate own-name processing. These studies examine whether there are differences at the neural level in how the own name is processed compared to other names. Research using fMRI or PET that contrasted the response to the auditorily presented own name with other names found activation in the medial frontal cortex and in the superior temporal cortex (Carmody & Lewis, 2006; Kampe et al., 2003; Perrin et al., 2005; Staffen et al., 2006). When controlling for familiarity, by contrasting the own name with that of a close other, the activity was specific to the right inferior frontal gyrus (Tacikowski et al., 2010). EEG studies using event-related potentials (ERPs) have found an enhanced late parietal positive deflection (referred to as the parietal P3, P3b or parietal positivity) for the own name compared to the names of (close) others (visual: Cygan et al., 2014; Folmer & Yingling, 1997; Tacikowski & Nowicka, 2010; auditory: Nijhof et al., 2018; Oomen et al., 2022). This parietal positive deflection occurs 300 ms or later after stimulus onset over central-parietal scalp electrodes and has consistently been associated with self-preferential processing and self-other distinction (Knyazev, 2013).

A limitation of the neuroscientific methods used so far is that they are highly susceptible to noise. That is, measurement noise from muscle contractions, head and eye movements and interference from other devices inevitably contaminate data (Norcia et al., 2015). To mitigate this limitation, long test sessions and large samples are required to gather robust data. This is however costly and not always feasible. In the past few years, researchers in social sciences and linguistics have created and validated tasks using EEG frequency tagging to resolve this issue of noise (e.g. social interaction recognition: Oomen et al., 2022, 2023; face processing: Rossion & Boremanse, 2011; lexical representation: Lochy et al., 2015). This technique uses the periodic repetition of stimuli to ‘tag’ the brain response confined to a small set of narrow frequency bins (Regan, 1966). For example, face stimuli presented two times per second (2 Hz) will induce a response in the EEG signal associated with face processing that is detectable in the frequency domain as a high amplitude ‘peak’ at exactly 2 Hz and its harmonic frequencies (integer multiples of the presentation frequency, i.e. 4 Hz, 6 Hz, etc.; Regan, 1966). As a result, the noise from other frequencies does not contaminate the narrow band response of interest, leading to a high signal-to-noise ratio (Retter & Rossion, 2016). This and the fact that no (long) interstimulus intervals are needed for baseline correction as typically applied in ERP research, allows for shorter task durations. Lastly, an advantage of frequency tagging is that the response of interest is defined objectively since it is coupled to the presentation frequency defined prior to data-collection (Norcia et al., 2015).

These advantages of the EEG frequency tagging method are especially relevant for research in populations that have difficulty sitting still or concentrating such as infants, young children, and certain clinical populations. At the same time, own-name processing is a particularly important topic in these groups, as the own name plays a key role in early social and cognitive development. For example, reduced responsiveness to one’s own name has been identified as one of the earliest markers of autism and a predictor of later diagnosis (e.g. Nadig et al., 2007)

The aim of this study was to create and validate an EEG frequency tagging task with which the self-preferential effect of the own name can be measured. Although most EEG frequency tagging studies to date have used visual stimuli, we opted for the auditory modality as the own name presented aurally does not only function as a self-relevant cue but also as a socially relevant (ostensive) cue, in contrast to a visually presented name. So far, auditory frequency tagging studies have been used with relatively simple stimuli, such as vowels (Bharadwaj et al., 2014), tones (Nozaradan et al., 2017), syllables and pseudowords (Pinto et al., 2022). Furthermore, a study by Barbero et al. (2021) investigated rapid automatic categorisation of spoken voice (including words) versus non-voice (e.g. musical instruments) stimuli. However, to our knowledge, no auditory frequency tagging study exists yet that has compared higher-level differential responses to different categories of words such as responses to different names (which are considered words, i.e., proper nouns). Similarly to the original behavioural dichotic listening task (Cherry, 1953; Moray, 1959), we chose to present the stimuli dichotically to investigate selective attention under direct competition, and to investigate brain lateralization for name processing (Hugdahl et al., 2009). Although there is generally a right-ear advantage in dichotic listening due to the left-hemispheric lateralisation of language (Kimura, 1961b, 1961a, 1967), the ear advantage can depend on the type of stimulus (Majidpour et al., 2022). In other words, this dichotic task could tell us in which hemisphere the processing of meaningful names (primarily) takes place. We therefore investigated if the brain response is influenced by the ear in which the own and other names are presented. To differentiate between a familiarity effect and self-preferential effect (see Amodeo et al., 2022), we included a stranger name and a close-other name condition besides the own name condition. More specifically, on each trial, participants heard two names simultaneously, one in each ear. One name was a filler name, and the other was one of the three experimental conditions – either the own name, a close-other name, or a recurring stranger name. We expected a self-preferential effect, that is: an enhanced response to the own name compared to the close-other name (and stranger name).

## Open science statement

Our hypotheses, study design and data analysis plan were pre-registered (study 1: https://aspredicted.org/DBW_2WF; study 2: https://aspredicted.org/BG6_4B3). Data and analyses can be found on the Open Science Framework (https://osf.io/gdcr4/?view_only=e84d1ef0bc624f9a8ab2912535d688ef)

## Study 1

### Methods

#### Participants

To determine the sample size, we conducted an a-priori power analysis (significance level: 0.05, power: 95%, effect size *d_z_:* .80). We expected a large effect size as a previous ERP auditory oddball study found a large effect for own versus close-other name processing (Nijhof et al., 2018). The power analysis showed that we needed at least 23 participants to detect such a large effect size. However, as this task was administered together with another unrelated task that required a sample of 33 to find a medium effect size, we tested 33 participants. In this way, we also accounted for a possible smaller effect size due to, for example, the difference in paradigm (dichotic listening versus oddball) and analysis technique (frequency tagging versus ERP) between this study and the study by Nijhof et al. (2018). All participants were 18 to 35 years old, right-handed, Dutch speaking, had normal hearing, normal or corrected-to-normal vision, no dreadlocks/cornrows, and no history of a neurological or psychiatric condition. Two participants were excluded due to bad data quality. Therefore, the final sample contained 31 participants (24 female 7 male, *M_age_* = 22.94, *SD* = 3.27). Participants were reimbursed for their time. The protocol of this study was approved by the local ethics committee of the Faculty of Psychology and Educational Sciences of Ghent University (EC/2018/23).

#### Procedure, stimuli, task

Prior to the day of test session, participants were asked to fill in a short form so we could individualize the task to each participant. Through this form, participants shared their first name and its phonetic pronunciation, along with the first name of a close other (e.g. a family member or a close friend) and its phonetic pronunciation. Additionally, they indicated from a list of 79 Flemish first names whether they knew someone with that name or not. On arrival to the test session, participants were seated in a Faraday cage ∼80–100 cm from a 24-inch computer monitor with a refresh rate of 60 Hz and signed an informed consent. During EEG preparation, all participants completed the Dutch version of the Autism Spectrum Quotient (AQ; Baron-Cohen et al., 2001; Dutch version: Hoekstra et al., 2010). The AQ had an acceptable internal consistency in the current sample (α = .78, *M* = 105.42, *SD* = 10.74^1^) and was included for exploratory purposes to relate potential self-preferential effects with autism traits, based on a previous study that found diminished own-name processing in autism (Nijhof et al., 2018). Next, the participants completed two tasks, one that is unrelated to the current study and therefore not described here. The task with which participants started was counterbalanced across participants. The entire test session lasted for approximately 1,5 hours including the preparation and removal of the EEG.

All name stimuli were recorded with a friendly intonation by the same two Dutch-speaking females. The recordings were normalized to have the same maximum volume using Audacity (https://www.audacityteam.org). The stimuli were presented binaurally through EEG-compatible insert earphones (ER-3C, MedCat). Mean and peak measurement of the sound level as measured over a single presentation block was LAeq = 70,5 dB(A) and LAFmax = 80,7 db (A), respectively. There were four types of stimuli: the participants’ own name, a close-other name, a recurring stranger name, and filler names. For each participant, the recurring stranger name and 48 filler names were randomly selected from the list of Flemish first names given to the participant prior to the day of the test session. Names that the participant indicated to associate with someone they knew were removed from this list, as were names that were similar to the own or close-other name. For example: if the participant’s name was Griet, we would remove the filler name Margriet from the list.

The task was programmed in PsychoPy (Peirce et al., 2019). On each trial, participants heard two names simultaneously at a frequency of 1 Hz, one in the left and one in the right ear. One of the names was always a filler name, whilst the other was one of three experimental names: the own name, close-other name, or the recurring stranger name. The experimental names were presented in random order to diminish predictability. The task included 5 blocks that each consisted of 96 trials (duration task: ∼ 7,5 mins; 1 block: 1,5 minutes). The 48 filler names were thus presented two times per block, once by each voice, in a random fashion. The experimental names were presented equally often to the left and right ear and by each voice. The two simultaneously presented stimuli were always spoken by the two different voices, as hearing the same voice say two different names simultaneously would be unnaturalistic.

Participant instructions were given via an instruction screen before the start of the task. To minimize eye movements, participants were instructed to focus their gaze on a cross positioned at the centre of the screen. The instruction screen also showed the experimental names. We informed them about these three recurring names to reduce the variability between the three experimental names regarding the timepoint at which participants recognized them. That is, the recurrence of the own name and the close-other name would otherwise most likely be noticed earlier than the recurring stranger name.

#### EEG recording and pre-processing

EEG was continuously recorded at a sampling rate of 1000 Hz with 64 active Ag/AgCI electrodes (ActiCAP, Munich, Germany), using an ActiCHamp amplifier and BrainVision Recorder software (version 1.21.04.02, Brain Products, Gilching, Germany). Electrodes were positioned according to the 10% system, except for two electrodes (TP9 and TP10) that were placed at OI1h and OI2h according to the 5% system to cover a wider area of posterior-occipital activation. This was needed for the other task administered during the test session that included visual stimuli. FT9 and FT10 electrodes were used to record horizontal EOG. Additional bipolar Ag/AgCl sintered ring electrodes were placed above and below the left eye to record vertical electrooculogram (EOG). Fpz was used as ground electrode and Fz as an online reference.

Off-line pre-processing of the raw data was done using Letswave 6. First, a fourth-order Butterworth band-pass filter was applied with a low and high cut-off of 0.1 Hz and 100 Hz. To remove eye blinks, we computed an ICA matrix for each participant using the Runica algorithm and a square matrix. For each participant, the first 10 independent components were inspected and those capturing eye blinks or horizontal eye movements were manually removed. Next, noisy electrodes were interpolated using three neighbouring electrodes (9 interpolations in total; max 2 per participant^2^). As the stimuli were presented in random order to diminish predictability, a false sequencing approach was used (see Quek & Rossion, 2017) to create a 160 s continuous EEG trace with a periodicity of exactly 1 Hz for each condition separately: the own name, close-other name, and the recurring stranger name. This was done by first segmenting according to the three conditions. Then we concatenated these 1000 ms segments of each condition by aligning segment X-1 and the first value of segment X. Finally, to correct for the introduced drift in the newly created ‘false sequences’ we applied another high pass filter (0.1 Hz). After false sequencing, the online reference (Fz) was included as a regular electrode and the data was re-referenced to an average reference. Next, a Fast Fourier Transform was applied to transform the data of each electrode to normalized (divided by N/2, where N is the length of the data) amplitudes (μV) in the frequency domain.

#### Statistical analysis

Frequency tagging elicits not only responses at the frequency of stimulation (F, 1 Hz here) but also across its higher harmonics (2F, 3F, etc.: 2, 3, etc. Hz here). To accurately capture the evoked brain response, we summed the baseline-subtracted amplitudes across all the relevant harmonics (Retter et al., 2021; Retter & Rossion, 2016). This was done as follows (for a similar approach see Cracco et al., 2022a, b): we first determined the number of harmonics to include by (i) taking the grand-averaged amplitudes across participants, conditions, and electrodes, (ii) calculating a z-score for each frequency bin using the average voltage amplitude of the 20 neighbouring bins as baseline (10 on each side, excluding the immediately adjacent bins), and (iii) identifying the harmonics with a z-score > 2.32. With this procedure we identified 8 harmonics (1 Hz to 8 Hz). For each of these harmonics we then calculated the baseline-subtracted amplitudes for every participant, condition, and electrode using the same baseline as for the z-scores (Norcia et al., 2015). This summed baseline-corrected response was used as the dependent variable.

To ensure an unbiased selection of electrodes, independent of condition effects or hypotheses, a collapsed localizer approach was used to determine the electrodes of interest (Luck & Gaspelin, 2017). More specifically, using the summed baseline-subtracted amplitudes we identified the relevant electrodes based on visual inspection of the averaged topography across participants and conditions (see Figure 1 for the collapsed topography). Inspection of this topography showed frontocentral activity (FC1, FCz, FC2), as well as lateralized temporoparietal activity (left: PO7, P7, TP7; right: PO8, P8, TP8).

**Figure 1.**
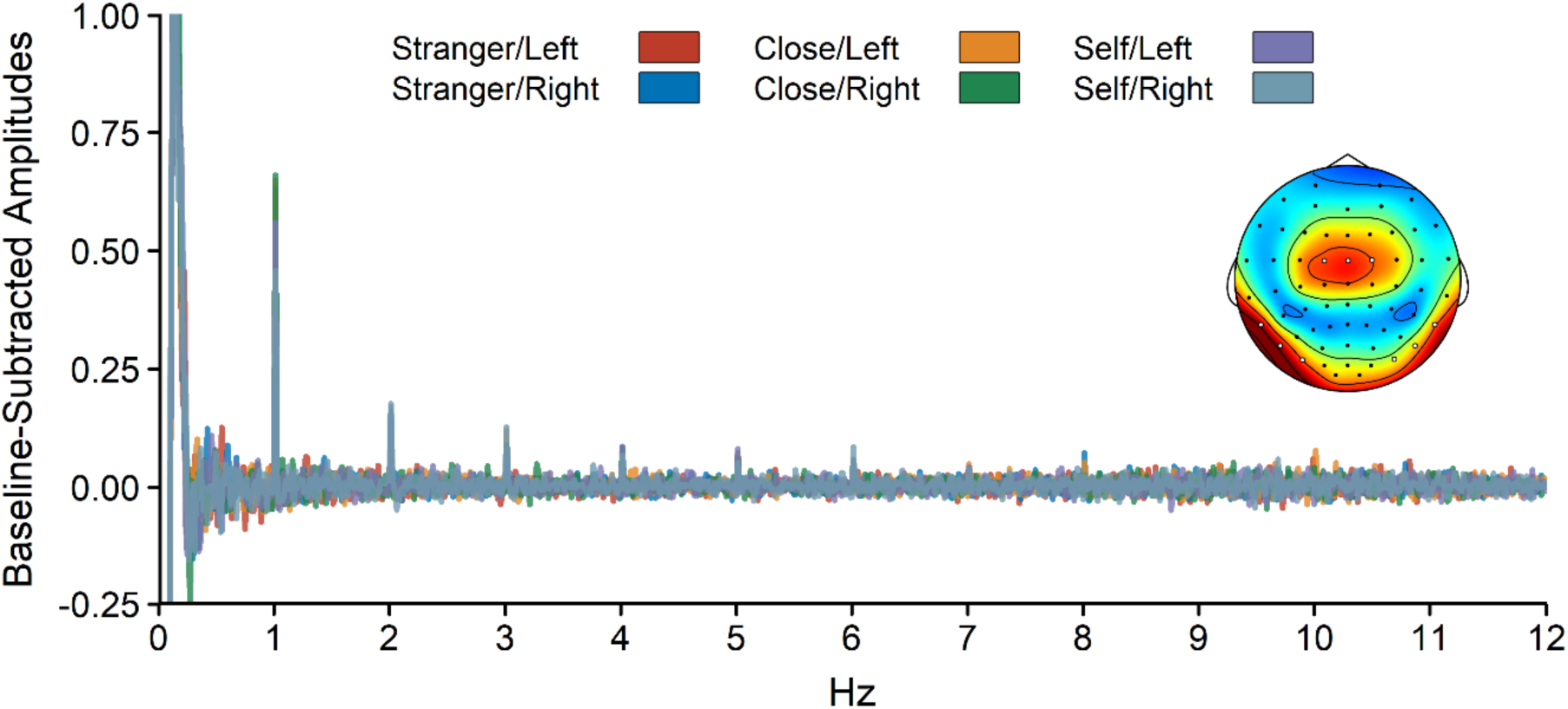
Baseline-subtracted amplitudes of the included electrodes separately for each name and ear and across participants, together with the collapsed topography (over participants and conditions). The topography is scaled from 0 to the maximum amplitude across all electrodes (i.e., 1.37 μV). Included electrodes are marked in white.

On the summed baseline-corrected data, we conducted a repeated measures MANOVA, with cluster (frontocentral, left temporoparietal, right temporoparietal), presentation side (left or right), and name (self, close other, or stranger) as within-subject factors. Significant interactions and main effects of cluster and name were followed up by two-tailed paired *t*-tests.

### Results

The repeated measures ANOVA (see boxplots and topographies per condition in Figure 2) revealed a main effect of cluster, *F*(2, 28) = 5.01, *p* = .014, η_p_^2^ = 0.26, with stronger responses in the left temporoparietal cluster than in the right temporoparietal cluster, *t*(29) = 3.00, *p* = .006, *d*_z_ = 0.55, and stronger responses in the frontocentral cluster than in the right temporoparietal cluster, *t*(29) = 2.95, *p* = .006, *d*_z_ = 0.54, but no difference between the left temporoparietal cluster and the frontocentral cluster, *t*(29) = 1.73, *p* = .095, *d*_z_ = 0.32. The effect of name was not significant, *F*(2, 28) = 1.89, *p* = .169, η_p_^2^ = 0.12, and, if anything, went in the opposite direction than expected, with responses being stronger to stranger and close other names than to the own name. None of the other effects were significant, all *F* ≤ 1.61, all *p* ≥ .203.

**Figure 2.**
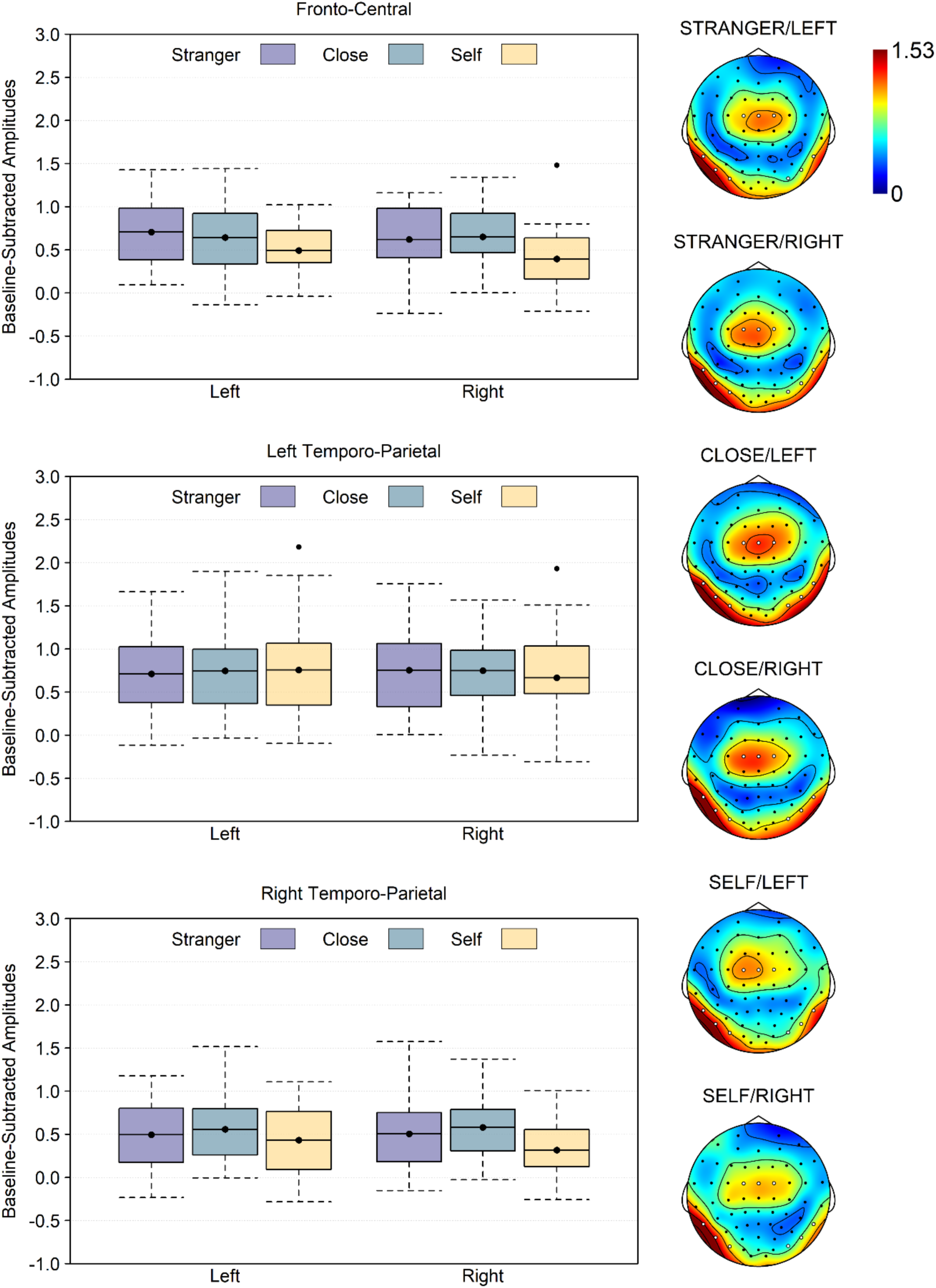
Left: Boxplots of the baseline-subtracted amplitudes per cluster, presentation side and name. Note that the boxplots show the mean instead of the median to match the statistical analysis, and that 0 is the baseline. As such, values below 0 necessarily reflect noise. Right: topographies per presentation side and name. Included electrodes are marked in white.

### Interim discussion

Previous research has shown that the own name is processed preferentially. Here, we investigated whether auditory EEG frequency tagging can be used to measure own-name processing, a technique that has the advantage of providing a high signal-to-noise ratio. Based on previous ERP research, we expected a self-preferential effect, that is, an enhanced response for the own name compared to the close-other and stranger name (Nijhof et al., 2018; Oomen et al., 2022). Additionally, we wanted to investigate the effect of the presentation ear on the processing of the names.

Contrary to our hypothesis, we did not find a significant effect of names. If anything, the opposite pattern to what was expected was observed, with descriptively (but non-significantly) stronger responses to close-other and stranger names than to the own name. We also did not find the presentation ear to influence the processing of the names. The only significant effect was for cluster. We found an overall stronger response at left temporoparietal and frontal central electrodes compared to right temporoparietal electrodes. This enhanced neural response in the left hemisphere is in line with studies that found left-hemispheric lateralisation of language (Kimura, 1961b, 1961a, 1967). The central activity found here is also typical for EEG studies that investigated name processing, particularly for later components, although this activity has previously been found more parietally (Nijhof et al., 2018; Oomen et al., 2022).

A possible reason for why we did not find a self-preferential effect might be related to the paradigm. That is, we chose to present the stimuli dichotically to investigate selective attention, similarly to the behavioural dichotic listening task (Cherry, 1953; Moray, 1959), but unlike other EEG studies that focused on name processing (Nijhof et al., 2018; Oomen et al., 2022). The opposite (non-significant) pattern that we found (stranger, close-other > own name) might be explained by this dichotic presentation style. To clarify, it might be that other names (the filler names presented to the other ear) get suppressed in favour of processing the own name, leading to an overall weaker total response. To confirm this post-hoc explanation, we ran a second study in which we presented the same name to both ears (i.e. non-dichotic). In line with study 1, we predicted a self-preferential effect for own name.

## Study 2

### Methods

The methods were nearly identical to that of study 1. In this Methods section of study 2 we only discuss deviations.

#### Participants

We tested the same number of participants as described in study 1, but with a different sample. One participant was excluded due to bad data quality, which resulted in a final sample of 32 participants (27 female, 5 male, *M*_age_ = 22.16, *SD* = 3.94).

#### Procedure, stimuli, task

The AQ had an excellent internal consistency in study 2 (α = .92, *M* = 101.50, *SD* = 18.60). The task was administered as the second task in a test session including three other unrelated tasks not further described here. Note that the task in study 2 included no filler names, as the experimental names were always presented in both ears simultaneously (non-dichotic). The names were presented randomly with one restriction: the same name spoken by the same voice was never repeated more than two times back-to-back.

#### Statistical analysis

Using the same procedure as in study 1, we identified 10 harmonics (1 Hz to 10 Hz) The averaged topography was similar to that of study 1, resulting in the same selection of electrodes and clusters (see Figure 3 for the collapsed topography). We performed a cluster (frontocentral, left temporoparietal, or right temporoparietal) x name (self, close-other, or stranger) repeated measures MANOVA.

**Figure 3.**
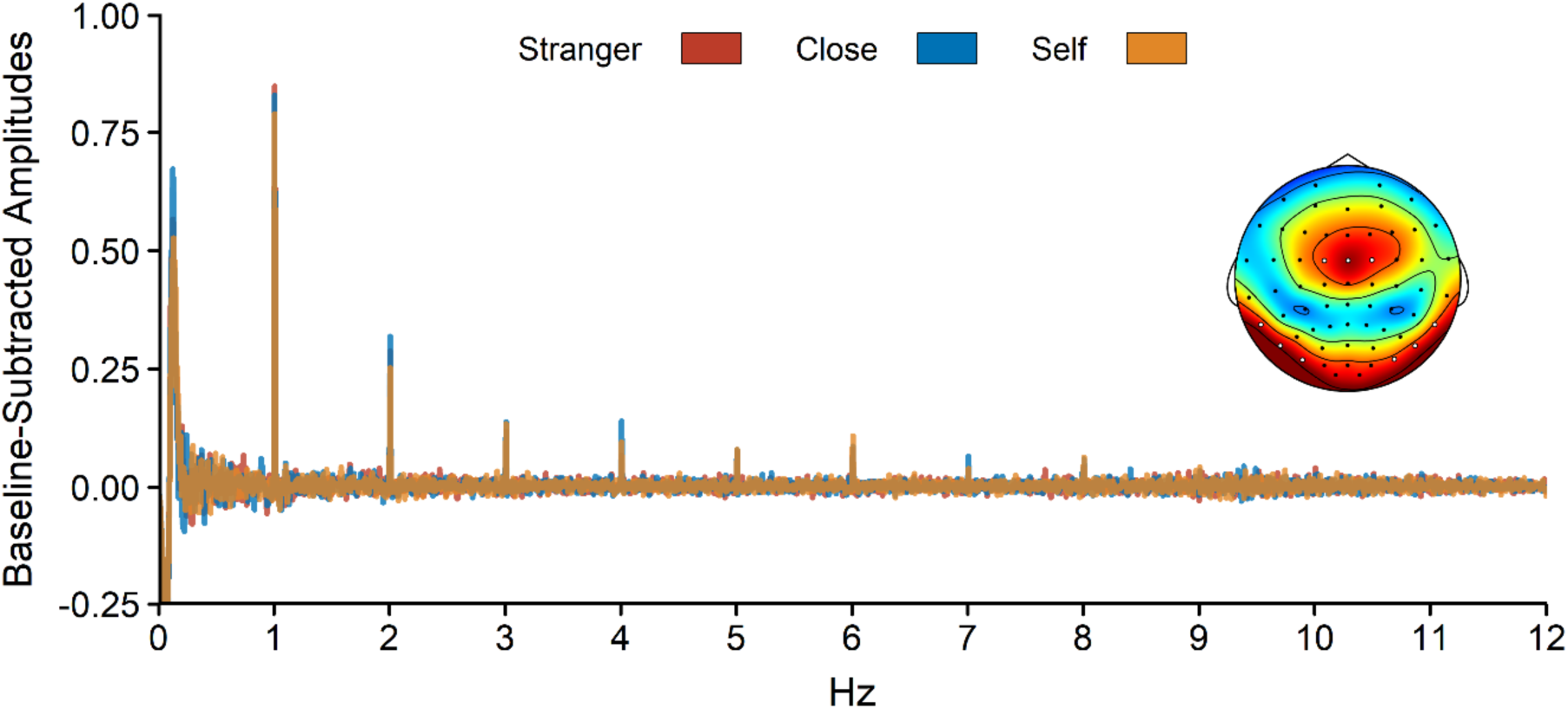
Baseline-subtracted amplitudes of the included electrodes separately for each name and across participants, together with the collapsed topography (over participants and conditions). The topography is scaled from 0 to the maximum amplitude across all electrodes (i.e., 1.84 μV). Included electrodes are marked in white.

### Results

The repeated measures ANOVA (see boxplots and topographies per condition in Figure 4) revealed no significant effects of cluster, *F*(2, 30) = 2.28, *p* = .120, η_p_^2^ = 0.13, name, *F*(2, 30) = 1.59, *p* = .221, η_p_^2^ = 0.10, or name x cluster, *F*(4, 28) = 0.93, *p* = .461, η_p_^2^ = 0.12.

**Figure 4.**
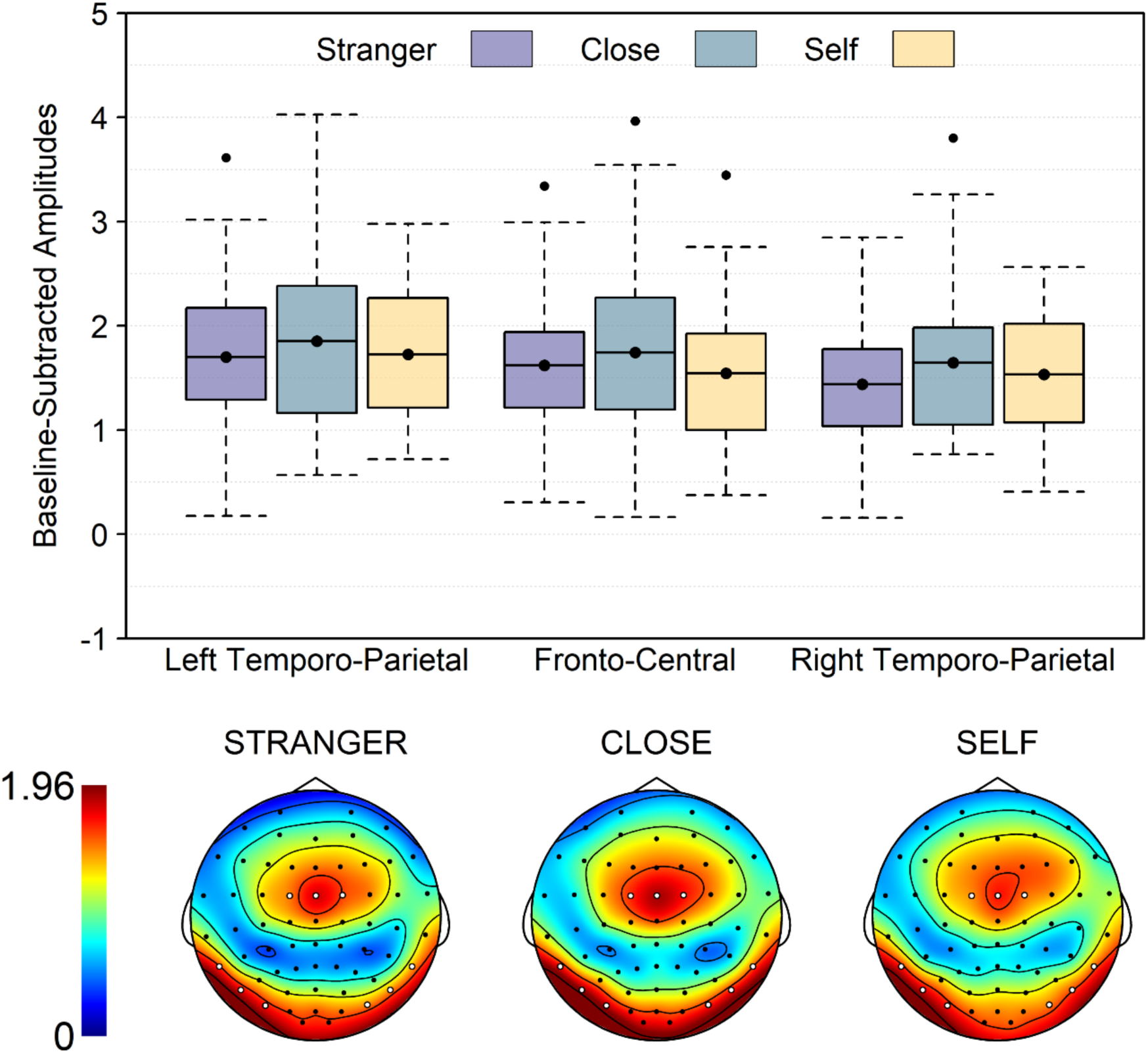
Top: Boxplots of the baseline-subtracted amplitudes per cluster and name. Note that the boxplots show the mean instead of the median to match the statistical analysis, and that 0 is the baseline. As such, values below 0 necessarily reflect noise. Bottom: Topographies per name. Included electrodes are marked in white.

### Interim discussion

Study 2 was conducted to investigate whether the dichotic design of study 1 led to the absence of a self-preferential effect. However, we did not find support for this explanation, as no influence of name was found in this study either. The question thus remains: why are we unable to capture the self-preferential effect as found by previous ERP studies (Nijhof et al., 2018; Oomen et al., 2022) with frequency tagging? A possible explanation could be that frequency tagging does not take temporal information into account. That is, in the frequency domain temporal information is collapsed, meaning it does not differentiate between early and late components of the response. This might be a problem for the specific effect we are interested in, as this self-preferential effect of names is typically associated with a late component, more specifically the amplification of a late parietal positive deflection (see Supplementary material for a visualization of the own-name effect found by Nijhof et al., 2018, in the time domain). We reasoned that the failure to find a self-preferential effect in our studies may be due to the fact that the effect of interest takes place at a late stage and that early brain responses may overshadow this later effect of interest. In a next step, we put this explanation to the test, by taking into account the timing of our effect of interest. More specifically, we did this by reanalyzing the data of study 1 and 2 but segmenting each trial into two parts: an early and late segment (500 ms each). If frequency tagging can capture the self-preferential effect, we would expect this effect to be present in the late segment.

#### Time-Resolved Frequency Tagging Analysis

To differentiate between early and late responses, each trial was segmented into an early and a late segment. This resulted in 500 ms segments instead of 1000 ms segments and thus a base frequency of 2 Hz instead of 1 Hz. Using the same procedure as in study 1, for the segmented study 1 data, we identified 4 harmonics for the early component and 1 for the late component. For the segmented study 2 data, we identified 7 for the early component and 3 for the late component. Further pre-processing steps were identical to those described in study 1.

#### Study 1

A segment x cluster x name repeated measures ANOVA revealed a main effect of cluster, *F*(2, 28) = 6.29, *p* = .006, η_p_^2^ = 0.31, with stronger responses in the left temporoparietal cluster than in the right temporoparietal cluster, *t*(29) = 3.63, *p* = .001, *d*_z_ = 0.66, and stronger responses in the frontocentral cluster than in the right temporoparietal cluster, *t*(29) = 3.38, *p* = .002, *d*_z_ = 0.62, but no difference between the left temporoparietal cluster and the frontocentral cluster, *t*(29) = 1.35, *p* = .187, *d*_z_ = 0.25. The main effect of segment was non-significant, *F*(1, 29) = 3.86, *p* = .059, η_p_^2^ = 0.12, but descriptively the data showed a stronger response in the early segment than in the late segment. The effects of name, *F*(2, 28) = 0.26, *p* = .776, η_p_^2^ = 0.02, segment x cluster, *F*(2, 28) = 0.63, *p* = .540, η_p_^2^ = 0.04, segment x name, *F*(2, 28) = 0.67, *p* = .520, η_p_^2^ = 0.05, cluster x name, *F*(4, 26) = 0.71, *p* = .592, η_p_^2^ = 0.10, and segment x cluster x name, *F*(4, 26) = 0.34, *p* = .851, η_p_^2^ = 0.05, were all non-significant.

#### Study 2

A segment x cluster x name repeated measures ANOVA revealed a main effect of segment, *F*(1, 31) = 133.66, *p* < .001, η_p_^2^ = 0.81, with stronger responses in the early than late segment, and a segment x name interaction, *F*(2, 30) = 5.46, *p* = .010, η_p_^2^ = 0.27, indicating that there was an effect of name for the late segment, *F*(2, 30) = 5.55, *p* = .009, η_p_^2^ = 0.27, but not for the early segment, *F*(2, 30) = 1.06, *p* = .360, η_p_^2^ = 0.07. The effect of name for the late segment indicated that responses were stronger to the own name than to both the close other, *t*(31) = 3.27, *p* = .003, *d*_z_ = 0.58, and the stranger name, *t*(31) = 2.87, *p* = .007, *d*_z_ = 0.51, whereas the response to the close other and stranger name did not differ, *t*(31) = 1.09, *p* = .283, *d_z_* = 0.19. In addition to these significant effects, there was a non-significant effect of cluster, *F*(2, 30) = 2.93, *p* = .069, η_p_^2^ = 0.16, and of segment x cluster, *F*(2, 30) = 3.27, *p* = .052, η_p_^2^ = 0.18, suggesting that responses to the left temporoparietal cluster and to the frontocentral cluster were stronger than responses to the right temporoparietal cluster in the early but not the late segment (see Figure 5). The main effect of name, *F*(2, 30) = 0.81, *p* = .453, η_p_^2^ = 0.05, the cluster x name interaction, *F*(2, 30) = 1.06, *p* = .394, η_p_^2^ = 0.13, and the segment x cluster x name interaction, *F*(4, 28) = 0.88, *p* = .490, η_p_^2^ = 0.11, were all non-significant.

**Figure 5.**
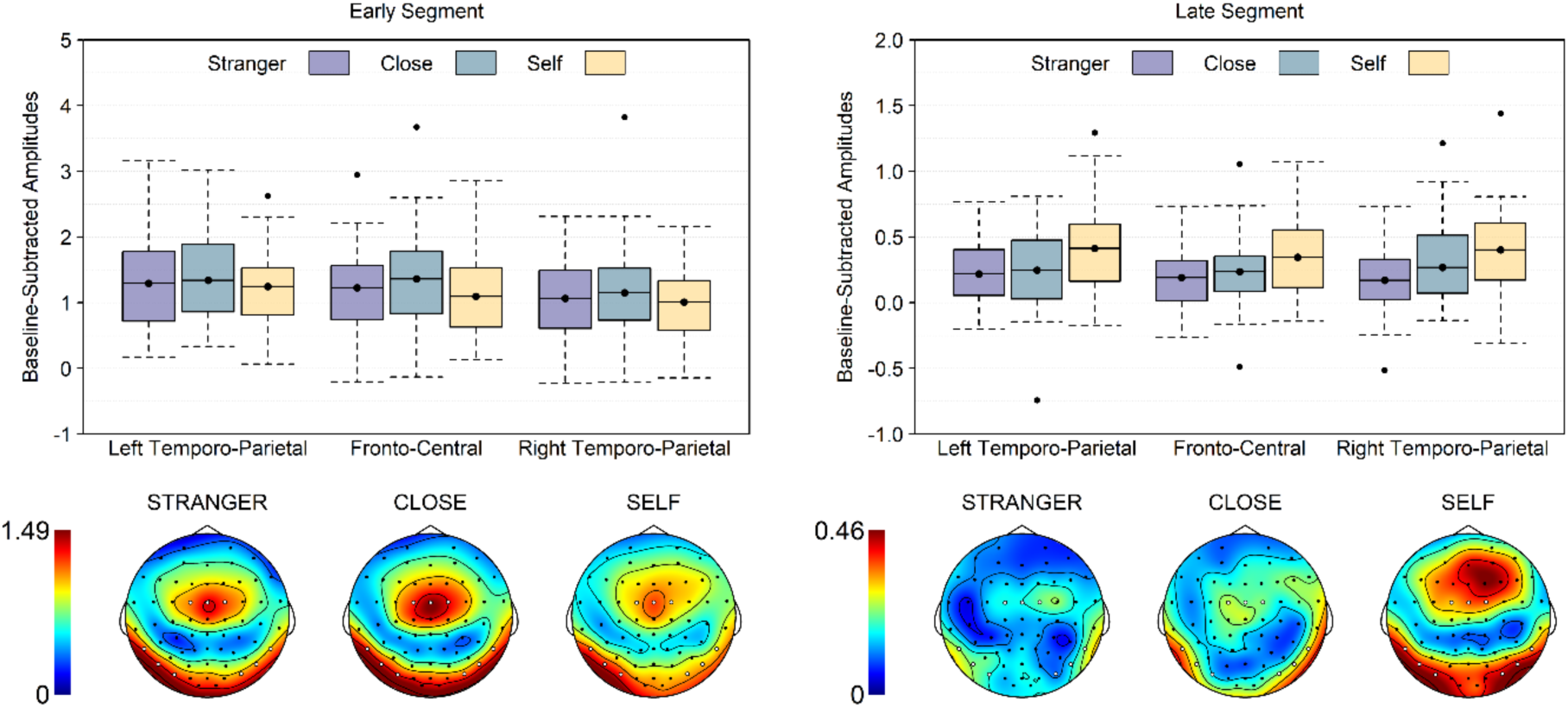
Top: Boxplots of the baseline-subtracted amplitudes per segment, cluster and name. Note that the boxplots show the mean instead of the median to match the statistical analysis, and that 0 is the baseline. As such, values below 0 necessarily reflect noise. Bottom: Topographies per name. Included electrodes are marked in white.

We also conducted an exploratory Spearman correlation to test whether own name processing (quantified as a difference index: own name – close-other name) relates to autism traits (AQ-scores). This analysis revealed no significant correlation, *r* = −0.06, *p* = .750).

### General discussion

The aim of this study was to create and validate an auditory EEG frequency tagging task to measure the self-preferential effect of one’s own name, leveraging its high signal-to-noise ratio (Norcia et al., 2015). In Study 1, we therefore created and tested an auditory frequency-tagging task in which name stimuli (own, close-other, stranger name) were presented dichotically to investigate selective attention under direct competition (similar to the cocktail party study design, e.g. Moray, 1959) and brain lateralization for name processing. However, there was no effect of either name or presentation ear.

We reasoned that the lack of a self-preferential effect may have been caused by using a dichotic listening paradigm, in contrast to the non-dichotic paradigms used in previous ERP research (e.g., Nijhof et al., 2018; Oomen et al., 2022). More specifically, as descriptive inspection of the data revealed an opposite pattern (stranger, close-other > own name) to what was expected, we suspected that the lack of an effect might be due to suppression of the filler names (the names presented to the opposite ear) in favor of processing the own name, leading to an overall weaker response. We addressed this post-hoc hypothesis in another experiment (Study 2) in which the stimuli were presented non-dichotically, but again found no effect of name.

We could thus rule out dichotic presentation as an explanation for our nonsignificant results. What could then be the reason? An obvious difference between ERPs and EEG frequency tagging is that the latter does not take into account the temporal information. Temporal information is collapsed, hence no differentiation is made between early and late components of the brain’s response to a stimulus. This is unlikely to pose a problem when investigating relatively simple perceptual processes, which are typically studied with EEG frequency tagging and reflected in early ERP components. However, it may have a crucial impact when studying more complex cognitive processes, such as self-preferential effects of the own name, which take place at later stages of information processing. We reasoned that collapsing temporal information may have obscured the self-preferential effect. That is, the effect of interest occurs late, but may have been overshadowed by early brain responses. To address this, we re-analyzed the data from both Study 1 and Study 2, making use of prior knowledge about the timing of the effect of interest. We refer to this approach as t*ime-resolved EEG frequency tagging.* Specifically, each trial was segmented into two 500 ms windows (early and late), with the expectation that the self-preferential effect would emerge only in the late window.

Reanalyzing the data from Study 1 (the dichotic task) did not reveal a significant interaction between name and segment, nor main effects of name or segment. This can likely be explained by the choice of the dichotic paradigm (deviating from previous ERP research) and by our original hypothesis of suppression of the filler names in favor of processing the experimental names. More importantly, re-analyzing the data from Study 2 (the non-dichotic task) revealed a self-preferential effect (own > close other, stranger name) specifically in the late segment, replicating previous ERP studies that reported a self-preferential effect at a late positive component (Cygan et al., 2014; Folmer & Yingling, 1997; Nijhof et al., 2018; Oomen et al., 2022; Tacikowski & Nowicka, 2010). No self-preferential effect was found in the early segment. However, the brain response was significantly stronger in the early than late segment, supporting our reasoning that the self-preferential effect may have been overshadowed by earlier brain activity.

The time-resolved EEG frequency tagging approach used in this study could be valuable for researchers using EEG frequency tagging who also want to study more complex cognitive processes. EEG frequency tagging as a method is not new, as steady-state visual evoked potentials (SSVEPs) were already reported by Adrian and Matthews (1934). Since then, however, the method has almost exclusively been used to study lower-level perceptual processes and attention, typically linked to early occurring ERP components in the time domain. Only recently have researchers begun to apply the method to study more complex cognitive processes (e.g., perspective taking: Beck et al., 2018; social interaction recognition: Oomen et al., 2022, 2023), demonstrating the suitability and added value of the method in these domains. Note, however, that studies investigating more complex cognitive processes may require lower frequencies (Oomen et al., 2022) to capture later appearing effects of interest, whereas traditional studies on low-level processes typically used frequencies in the 3-20 Hz range, suited for early ERP components. Our findings point to a potential second pitfall of using EEG frequency tagging for studying more complex cognitive processes: collapsing temporal information can obscure later effects. The time-resolved EEG frequency tagging approach addresses this by leveraging temporal-domain knowledge, providing a tool that other researchers can apply when studying complex cognitive processes.

Perhaps the most obvious example of a study that could benefit from the time-resolved approach is the study by Nijhof et al. (2024). They found a self-preferential effect in their EEG frequency study for own faces but not for (written) own names, which contrasts with existing ERP research that showed a self-preferential effect both for own name (Tacikowski & Nowicka, 2010; Tacikowski et al., 2011) and own faces (Alzueta et al., 2019; Oomen et al., 2022). However, the self-preferential effect for written own name was found to be reflected in the modulation of a late parietal positive deflection, whereas the self-preferential effect for faces has been shown to modulate an early ERP component (N250, Alzueta et al., 2019; Oomen et al., 2022), which may explain why Nijhof et al. (2024) were able to find a self-preferential effect for faces but not names in their study, for similar reasons as in our study.

To conclude, we created and validated an auditory EEG frequency tagging task with which to measure the self-referential effect of the own-name and replicated previous ERP studies that found stronger neural responses for the own name compared to close-other names. This task is particularly useful for studies where large samples or long experiments are not feasible (infants and clinical populations such as individuals with autism), given its high signal-to-noise ratio. Furthermore, the time-resolved EEG frequency tagging approach used here can be used in future studies on other complex processes that are associated with late components.

## Acknowledgement

DO was supported by the Special Research Fund of Ghent University (BOF18/DOC/348). EC was supported by a postdoctoral fellowship awarded by the Research Foundation Flanders (12U0322N).

## Supplementary material

**Supplementary Figure 1.**
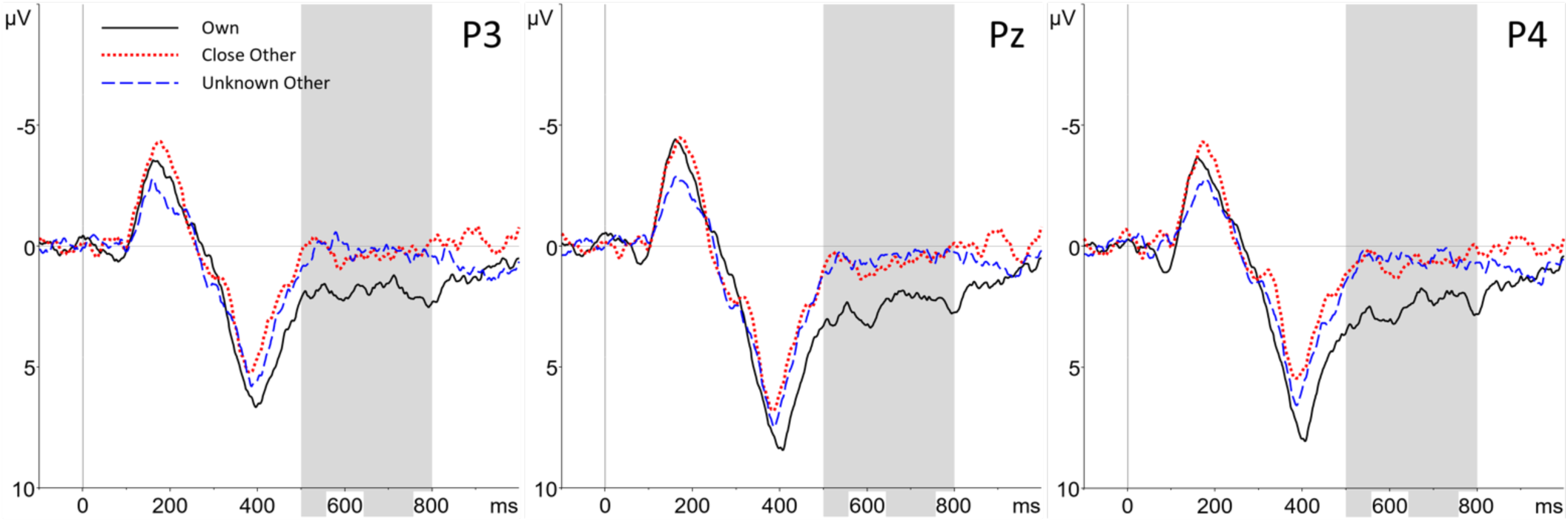
Grand average waveforms for the three electrodes (P3, Pz and P4) included in the parietal positivity analysis of Nijhof et al., 2018, revealing the self-preferential effect.

## Footnotes

1 Note that these statistics were calculated following the procedure of Hoekstra et al. (2010), who use the original 4-point Likert scale scores, unlike the procedure of Baron-Cohen et al. (2001) who dichotomize the answer categories.

2 The two participants excluded due to bad data quality would have required more than 10% of their electrodes to be interpolated.

## References

Alexopoulos, T., Muller, D., Ric, F., & Marendaz, C. (2012). I, me, mine: Automatic attentional capture by self-related stimuli. European Journal of Social Psychology, 42(6), 770–779. 10.1002/EJSP.1882

American Psychiatric Association. (2013). DSM-5 Diagnostic Classification. In Diagnostic and Statistical Manual of Mental Disorders. 10.1176/appi.books.9780890425596.x00diagnosticclassification

Arnell, K. M., Shapiro, K. L., & Sorensen, R. E. (2010). Reduced Repetition Blindness for One’s Own Name. 10.1080/135062899394876, 6(6), 609–635. https://doi.org/10.1080/135062899394876

Barbero, F. M., Calce, R. P., Talwar, S., Rossion, B., & Collignon, O. (2021). Fast Periodic Auditory Stimulation Reveals a Robust Categorical Response to Voices in the Human Brain. ENeuro, 8(3). 10.1523/ENEURO.0471-20.2021

Bharadwaj, H. M., Lee, A. K. C., Shinn-Cunningham, B. G., Ciaramitaro, V., & Shinn, B. G. (2014). Measuring auditory selective attention using frequency tagging. 10.3389/fnint.2014.00006

Carmody, D. P., & Lewis, M. (2006). Brain activation when hearing one’s own and others’ names. Brain Research, 1116(1), 153–158. 10.1016/J.BRAINRES.2006.07.121

Cherry, E. C. (1953). Some Experiments on the Recognition of Speech, with One and with Two Ears. The Journal of the Acoustical Society of America, 25(5), 975–979. 10.1121/1.1907229

Cygan, H. B., Tacikowski, P., Ostaszewski, P., Chojnicka, I., & Nowicka, A. (2014). Neural correlates of own name and own face detection in autism spectrum disorder. PLoS ONE, 9(1). 10.1371/journal.pone.0086020

Folmer, R. L., & Yingling, C. D. (1997). Auditory P3 responses to name stimuli. Brain and Language, 56(2), 306–311. 10.1006/brln.1997.1828

Hugdahl, K., Westerhausen, R., Alho, K., Medvedev, S., Laine, M., & HÄmÄläinen, H. (2009). Attention and cognitive control: Unfolding the dichotic listening story. Scandinavian Journal of Psychology, 50(1), 11–22. 10.1111/J.1467-9450.2008.00676.X

Kamp, A., Sem-Jacobsen, C. W., Leeuwen, W. S. van, & T-Weel, L. H. van der. (1960). Cortical responses to modulated light in the human subject. Acta Physiologica Scandinavica, 48(1), 1–12. 10.1111/j.1748-1716.1960.tb01840.x

Kampe, K. K. W., Frith, C. D., & Frith, U. (2003). “Hey John”: signals conveying communicative intention toward the self activate brain regions associated with “mentalizing,” regardless of modality. The Journal of Neuroscience : The Official Journal of the Society for Neuroscience, 23(12), 5258–5263. 10.1523/JNEUROSCI.23-12-05258.2003

Kim, K., Johnson, J. D., Rothschild, D. J., & Johnson, M. K. (2019). Merely presenting one’s own name along with target items is insufficient to produce a memory advantage for the items: A critical role of relational processing. Psychonomic Bulletin and Review, 26(1), 360–366. 10.3758/S13423-018-1515-9/FIGURES/1

Kimura, D. (1961a). Cerebral dominance and the perception of verbal stimuli. Canadian Journal of Psychology / Revue Canadienne de Psychologie, 15(3), 166–171. 10.1037/H0083219

Kimura, D. (1961b). Some effects of temporal-lobe damage on auditory perception. Canadian Journal of Psychology, 15, 156–165. 10.1037/H0083218

Kimura, D. (1967). Functional Asymmetry of the Brain in Dichotic Listening. Cortex, 3(2), 163–178. 10.1016/S0010-9452(67)80010-8

Knyazev, G. G. (2013). EEG correlates of self-referential processing. Frontiers in human neuroscience, 7, 264.

Lachter, J., Forster, K. I., & Ruthruff, E. (2004). Forty-five years after Broadbent (1958): still no identification without attention. Psychological Review, 111(4), 880–913. 10.1037/0033-295X.111.4.880

Lochy, A., Van Belle, G., & Rossion, B. (2015). A robust index of lexical representation in the left occipito-temporal cortex as evidenced by EEG responses to fast periodic visual stimulation. Neuropsychologia, 66, 18–31. 10.1016/J.NEUROPSYCHOLOGIA.2014.11.007

Majidpour, A., Aleaba, M. M., Aghamolaei, M., & Nazeri, A. (2022). Review of the Factors Affecting Dichotic Listening. Auditory and Vestibular Research, 31(2), 74–83. 10.18502/AVR.V31I2.9111

Moray, N. (1959). Attention in Dichotic Listening: Affective Cues and the Influence of Instructions, 11(1), 56–60. 10.1080/17470215908416289

Nijhof, A. D., Catmur, C., Brewer, R., Coll, M. P., Wiersema, J. R. & Bird, G. (2024). Differences in own-face but not own-name discrimination between autistic and neurotypical adults: A fast periodic visual stimulation-EEG study. Cortex, 171, 308–318. 10.1016/j.cortex.2023.10.023

Nijhof, A. D., Dhar, M., Goris, J., Brass, M., & Wiersema, J. R. (2018). Atypical neural responding to hearing one’s own name in adults with ASD. Journal of Abnormal Psychology, 127(1), 129–138. 10.1037/abn0000329

Nijhof, A. D., von Trott zu Solz, J., Catmur, C., & Bird, G. (2021). Equivalent own name bias in autism: An EEG study of the Attentional Blink. Cognitive, affective & behavioral neuroscience, 22(3), 625–639. 10.3758/s13415-021-009670-w

Norcia, A. M., Gregory Appelbaum, L., Ales, J. M., Cottereau, B. R., & Rossion, B. (2015). The steady-state visual evoked potential in vision research: A review. Journal of Vision, 15(6), 1–46. 10.1167/15.6.4

Nozaradan, S., Mouraux, A., & Cousineau, M. (2017). Sensory Processing: Frequency tagging to track the neural processing of contrast in fast, continuous sound sequences. Journal of Neurophysiology, 118(1), 243. 10.1152/JN.00971.2016

Oomen, D., Cracco, E., Brass, M., & Wiersema, J. R. (2022). EEG frequency tagging evidence of social interaction recognition. Social Cognitive and Affective Neuroscience, 00, 1–10. 10.1093/SCAN/NSAC032

Oomen, D., Cracco, E., Brass, M., & Wiersema, J. R. (2023). Frequency tagging evidence of intact social interaction recognition in adults with autism. Autism Research, 16(6), 1111–1123. 10.1002/aur.2929

Oomen, D., El Kaddouri, R., Brass, M., & Wiersema, J. R. (2022). Neural correlates of own name and own face processing in neurotypical adults scoring low versus high on symptomatology of autism spectrum disorder. Biological Psychology, 172, 108358. 10.1016/J.BIOPSYCHO.2022.108358

Perrin, F., Maquet, P., Peigneux, P., Ruby, P., Degueldre, C., Balteau, E., Del Fiore, G., Moonen, G., Luxen, A., & Laureys, S. (2005). Neural mechanisms involved in the detection of our first name: a combined ERPs and PET study. Neuropsychologia, 43(1), 12–19. 10.1016/J.NEUROPSYCHOLOGIA.2004.07.002

Pinto, D., Prior, A., & Golumbic, E. Z. (2022). Assessing the Sensitivity of EEG-Based Frequency-Tagging as a Metric for Statistical Learning. Neurobiology of Language, 3(2), 214–234. 10.1162/NOL_A_00061

Polich, J. (2007). Updating P300: An Integrative Theory of P3a and P3b. Clinical Neurophysiology : Official Journal of the International Federation of Clinical Neurophysiology, 118(10), 2128. 10.1016/J.CLINPH.2007.04.019

Regan, D. (1966). Some characteristics of average steady-state and transient responses evoked by modulated light. Electroencephalography and Clinical Neurophysiology, 20(3), 238–248. 10.1016/0013-4694(66)90088-5

Retter, T. L., & Rossion, B. (2016). Uncovering the neural magnitude and spatio-temporal dynamics of natural image categorization in a fast visual stream. Neuropsychologia, 91, 9–28. 10.1016/J.NEUROPSYCHOLOGIA.2016.07.028

Rossion, B., & Boremanse, A. (2011). Robust sensitivity to facial identity in the right human occipito-temporal cortex as revealed by steady-state visual-evoked potentials. Journal of Vision, 11(2), 16–16. 10.1167/11.2.16

Shapiro, K. L., Caldwell, J., & Sorensen, R. E. (1997). Personal names and the attentional blink: A visual “cocktail party” effect. Journal of Experimental Psychology: Human Perception and Performance, 23(2), 504–514. 10.1037//0096-1523.23.2.504

Staffen, W., Kronbichler, M., Aichhorn, M., Mair, A., & Ladurner, G. (2006). Selective brain activity in response to one’s own name in the persistent vegetative state. Journal of Neurology, Neurosurgery, and Psychiatry, 77(12), 1383. 10.1136/JNNP.2006.095166

Tacikowski, P., Brechmann, A., Marchewka, A., Jednoróg, K., Dobrowolny, M., & Nowicka, A. (2010). Is it about the self or the significance? An fMRI study of self-name recognition. 10.1080/17470919.2010.490665, 6(1), 98–107. https://doi.org/10.1080/17470919.2010.490665

Tacikowski, P., & Nowicka, A. (2010). Allocation of attention to self-name and self-face: An ERP study. Biological Psychology, 84(2), 318–324. 10.1016/j.biopsycho.2010.03.009

Woodman G. F. (2010). A brief introduction to the use of event-related potentials in studies of perception and attention. Attention, perception & psychophysics, 72(8), 2031–2046. 10.3758/APP.72.8.2031

